# Valence of sensory stimulation as a key feature for subcortical entrainment: Insights from human intracranial EEG

**DOI:** 10.1101/2025.01.03.631218

**Authors:** Roxane S. Hoyer, Marie-Frédérique Bernier, Laurence Martineau, Paule Lessard Bonaventure, Corentin Labelle, Arthur Borderie, Isabelle Blanchette, Anna Blumenthal, Steven Laureys, Philippe Albouy

**Author notes:** corresponding author, Roxane S. Hoyer.

## Abstract

Sensory rhythmic stimulation enhances executive functions by entraining oscillations in higher-order cortical networks, but its effects on subcortical structures remain unclear. We propose that stimulus valence is a key feature to enable subcortical entrainment. Using intracranial EEG in epileptic patients, we first show that visual search is supported by cortico-subcortical theta (5Hz) activity. We then show that 5 Hz negative-valence visual stimulation entrains theta oscillations in a task-related network, including the ventral visual stream, hippocampus, and dorsolateral prefrontal cortex. Finally, in a behavioral experiment in healthy individuals, we show that both neutral and negative valence 5 Hz stimulation improved visual search speed, but only negative valence stimulation enhanced target image recognition as assessed through an additional memory task. These findings highlight the role of stimulus valence in modulating subcortical brain activity and behaviors through rhythmic sensory stimulation and pave the way for further applications in clinical intervention.

## Introduction

Brain activity has the remarkable property to mimic the rhythm of environmental sensory (e.g., visual, auditory) information^1–9^. Oscillatory activity synchronizes to external rhythms^3,8^ via phase-locking^2^. This entrainment is not only observable in sensory cortices^5,6,9^, with possible cross-modal effects^2,9^, but also propagates in a supra-modal way through higher-order brain areas supporting cognitive functions such as attention^3–7,10,11^ and memory^2^. However, neuroscience research on oscillatory entrainment in higher-order brain networks remains an emerging area, requiring further development to define optimal stimulation parameters that facilitate supra-modal entrainment^2,12^. Particularly, while entrainment in cortical higher-order networks is currently gaining increasing interest^2,10,11^, it remains unknown whether subcortical activity can be entrained through external sensory rhythms and whether this entrainment can shape cognitive performance. Most studies, however, have employed neutral, non-emotional stimuli for rhythmic stimulation, leaving open the question of whether emotional valence can extend or intensify entrainment, particularly through the recruitment of subcortical circuits implicated in emotion and memory.

Given that negative emotional content robustly engages limbic circuits, including the hippocampus^13–15^, we hypothesized that valence-specific rhythmic stimulation would effectively recruit both cortical and subcortical networks at the stimulation frequency, thereby enhancing attentional processes and memory encoding beyond what can be observed with neutral stimuli. Consequently, we predicted that negative-valence stimulation would yield deeper entrainment and results in stronger higher order brain functioning and behavioral facilitation.

We used SEEG recordings in six patients with intractable epilepsy (see Supplemental Information [SI] S1) to examine cortical and subcortical entrainment by valence-specific (neutral and negative) rhythmic visual stimulation (RVS). Although the SEEG sample is necessarily small due to clinical constraints, this approach grants unparalleled spatiotemporal access subcortical brain activity – for instance in the hippocampus – to assess subcortical entrainment. Subsequently, we evaluated the effects of RVS on attention and memory encoding in a behavioral experiment performed in 27 healthy young adults.

By studying whether emotional stimuli can drive deeper neural synchronization, this study aims to inform both theoretical models of rhythmic sensory entrainment and future clinical non-invasive brain stimulation protocol leveraging emotional salience as a key stimulation parameter.

## Results

### Visual search is associated with sustained increased theta and gamma power in the occipital cortex

SEEG patients performed a visual search task. During visual search trials, they had to identify whether four grey-shaded items all came from the same category or one differed (e.g., 3 submarines and 1 whale; Method). All task blocks included equal proportions of No-Target (e.g., 4 submarines), Neutral-Target (e.g., 3 submarines and 1 whale), and Threatening-Target (e.g., 3 submarines and 1 shark) trials. Both living and non-living items were used as targets (Fig. 1.A, note that all patients performed the task above chance, see SI S2).

**Figure 1.**
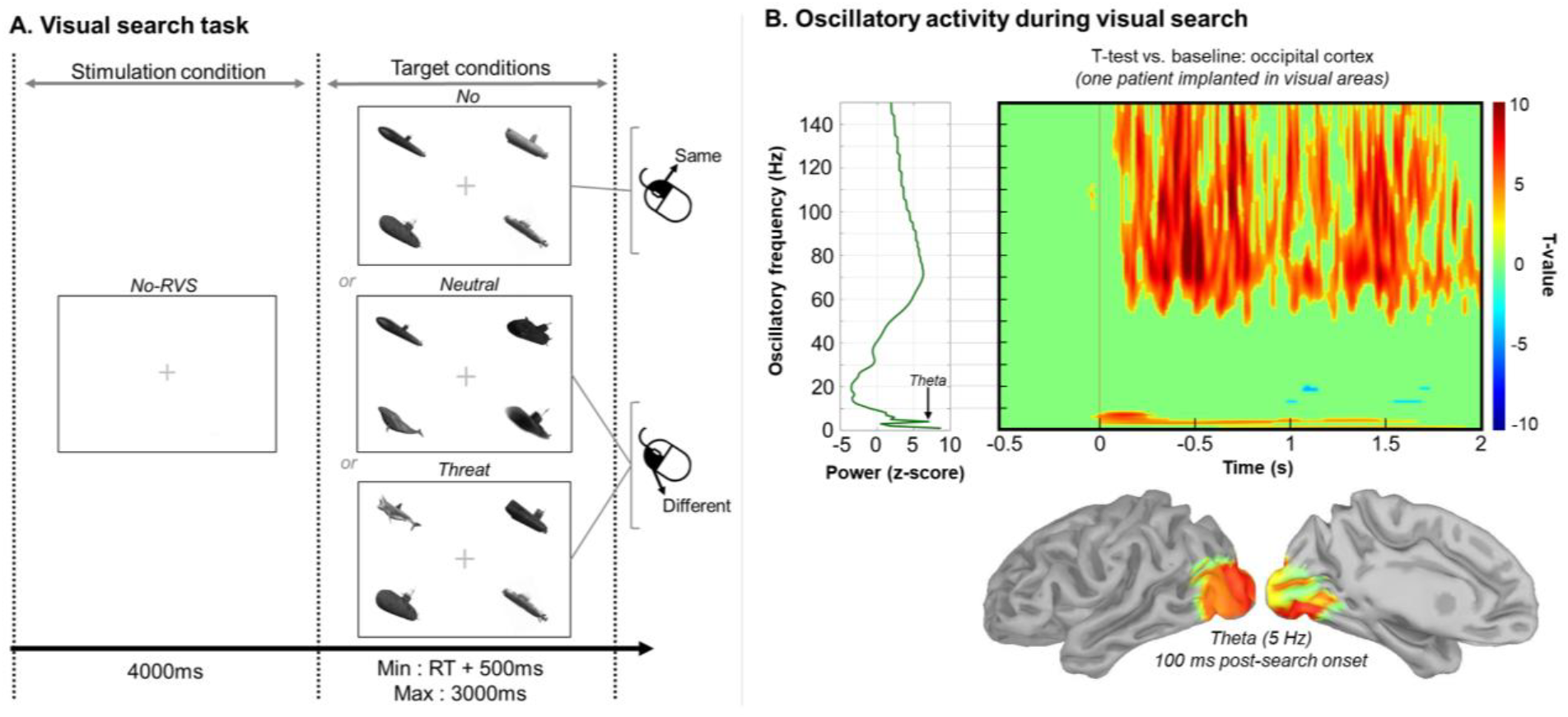
A) During visual search trials, participants decided if four grey-shaded items were from the same category or if one differed by pressing the right or left mouse button. Equal numbers of No-Target, Neutral-Target, and Threat- Target trials were presented. B) Left Panel: Magnitude (z-scored with baseline activity) for 0-2s period relative to stimulus onset across frequencies for SEEG contacts located in the occipital cortex. Right Panel: T values in the time frequency domain (t-test vs. baseline, FDR corrected, *p* < .001) for oscillatory activity during visual search for SEEG contacts located in the occipital cortex. The time frequency map indicates sustained theta (4-8 Hz) and gamma (60-120 Hz) activity across inferior, middle, and superior occipital gyri.

We first investigated the sensory response associated with the visual search. Figure 1.B shows t-values in the time-frequency domain across trials of the visual search task (Morlet decomposition^16^, False Discovery Rate[FDR]-corrected *t*-test vs. baseline, *p* < .001) for SEEG contacts located in the right visual cortex (AAL3 atlas, -0.5 to 2 s relative to stimulus onset, Method). This analysis revealed that the magnitude of theta (4-8 Hz) activity increases post- search onset (0-300 ms period) compared to baseline, while both theta and gamma (60-120 Hz) activity persisted until target detection.

### Cortico-subcortical theta power supports cognitive engagement during visual search

We next aimed to identify how theta activity propagates to brain networks to support visual search, by focusing on theta power activity (identified in Fig. 1), a well-established marker of inter-regional communication during cognitive tasks^17,18^. We extracted Hilbert transformed magnitude of the theta band (4-8 Hz, depicted in Fig. 1.B baseline corrected with z-score) during the visual search task and during a separate Rest condition for each SEEG contact and each trial. Significant differences between task engagement and resting-state were evaluated for each subject using two-tail paired *t*-tests with Monte Carlo simulations (1000) and FDR correction (all *ps* < .01) across time. To ensure the robustness of the results for identifying task-related brain network, we further employed a spatial cluster-based approach by keeping only overlapping SEEG contacts across electrodes or subjects (see Method and Borderie et al., 2024^19^, see also SI Tab. S3). During visual search, both early and transient (attention orienting and recognition) and late and sustained (higher-order and predictive mechanisms) processes are assumed to be involved in task execution^20–23^. Therefore, we used time series clustering approach (Dynamic Time Warping and Kmedoids, Method) to objectively isolate theta patterns emerging during visual search (only for significant SEEG contacts, see SI Fig. S4). This analysis was performed on the difference in theta magnitude during task performance and Rest condition, over a period ranging from 0 and 1.5 s relative to search onset.

As expected, spatial and temporal clustering revealed transient and sustained theta fluctuations during visual search (Fig. 2, purple and blue line, respectively)^20–23^. Search-onset elicited a transient theta increase followed by a late, and slightly longer response in brain regions listed in Fig. 2 (in blue, left panel). In parallel, theta power progressively increased in the amygdala, the hippocampus, the anterior insula and in the dorsolateral prefrontal cortex (DLPFC).Once these task-related cortico-subcortical networks had been defined, we then questioned in the same SEEG patients whether those distributed brain areas could be entrained by rhythmic sensory stimulations, with the hypothesis that varying the valence of the rhythmic stimulus will enable sub-cortical entrainment.

**Figure 2.**
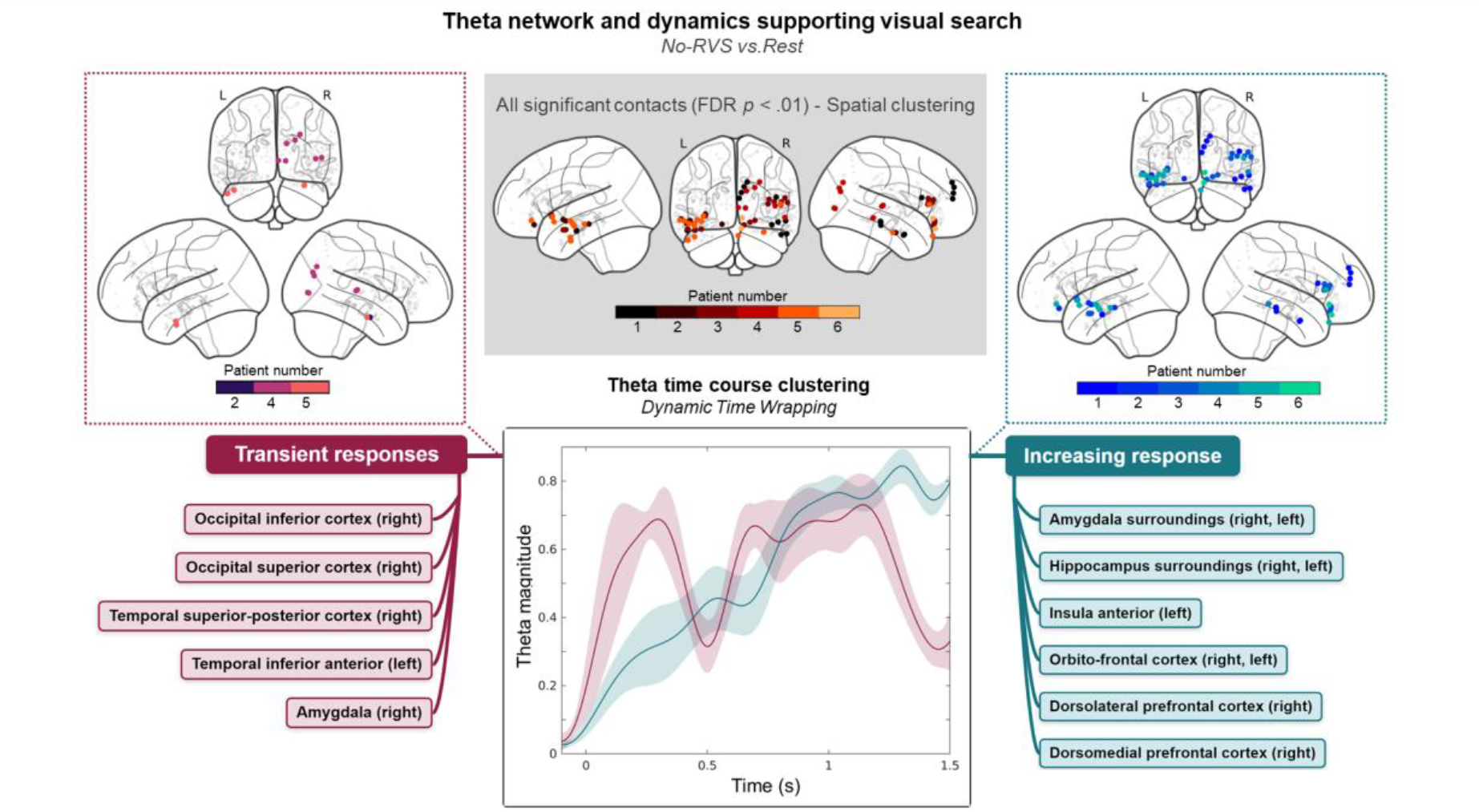
Spatial distribution and time courses of theta activity during visual search (−100 to 1500 ms). Middle panel: the top middle panel shows all significant SEEG contacts for the contrast task vs. rest across participants (clusters of at least two overlapping SEEG contacts). The lower middle panels show the time courses of theta activity for i) early evoked activity and later transient (purple) increase of theta oscillations for SEEG contacts located in occipital areas, the visual ventral and dorsal pathways, and the amygdala (left panel); ii) Sustained and increasing theta activity during visual search (blue line) for SEEG contacts located (MNI space) close to the amygdala and the hippocampus, the anterior insula, the orbito-frontal cortex, and the dorsolateral and dorsomedial prefrontal cortices (right panel). Shaded error bars indicate the standard error of the mean (SEM).

### Theta rhythmic visual stimulations elicit entrainment in task related cortical and subcortical networks

We studied the entrainment of oscillatory neurophysiological activity induced by 5 Hz flickering visual stimuli. The choice of 5 Hz, in the theta band, was made a priori on the basis of converging evidence that theta-frequency stimulation selectively engages the fronto-parietal (i.e., central-executive) network has been show to facilitates both attentional orienting and working-memory maintenance^2,12,17,18,22^. Before each trial of the visual search task, we presented 5 Hz RVS (4 s stimulation) to synchronize brain oscillatory activity and prime the task network and investigated selective entrainment by varying stimulus valence (Negative- RVS, snake stimulation; Neutral-RVS, rope stimulation – stimuli not used as target in the visual search task).

To evaluate neural entrainment, we computed the stimulation-to-brain phase-locking value (Stim-Brain PLV^24^) and investigated the phase relationship between the patients’ 5 Hz ongoing oscillations and a theoretical pure 5 Hz sinusoid (reflecting rhythmic visual stimulation) over the trial time window. The task period considered ranged from -4 to 0 s (stimulation period) and 0 to 2 s post-search onset (response period). This analysis was done for each trial, each SEEG contact, and for all participants. We then contrasted, for each patient, No-RVS, Neutral- RVS and Negative -RVS (Negative-RVS) using two-tail paired t-tests (*p* < .05) with Monte Carlo simulations (1000). To identify networks entrained with the 5 Hz RVS at the group level, a cluster-based method was employed: only groups of adjacent SEEG contacts (minimum: 2) exhibiting statistically significant increased Stim-Brain PLV were considered^19^ (see SI S5).

This analysis revealed that PLV was significantly higher in Negative-RVS compared to both No- and Neutral-RVS during the stimulation period. In the Negative-RVS compared to No- RVS, the Stim-Brain PLV was specifically enhanced in the inferior occipital cortex (right), medial temporal gyrus (left), hippocampus and its lateral surroundings (right), anterior insula (right), and DLPFC (right, Fig. 3.B and SI S3 and S4). In Negative-RVS compared to Neutral- RVS, the PLV was increased in the medial temporal gyrus (left), hippocampus and its surroundings, and DLPFC (right, Fig. 3.C and SI Fig. S3). No significant differences between stimulation conditions were observed during the response period after cluster correction.

**Figure 3.**
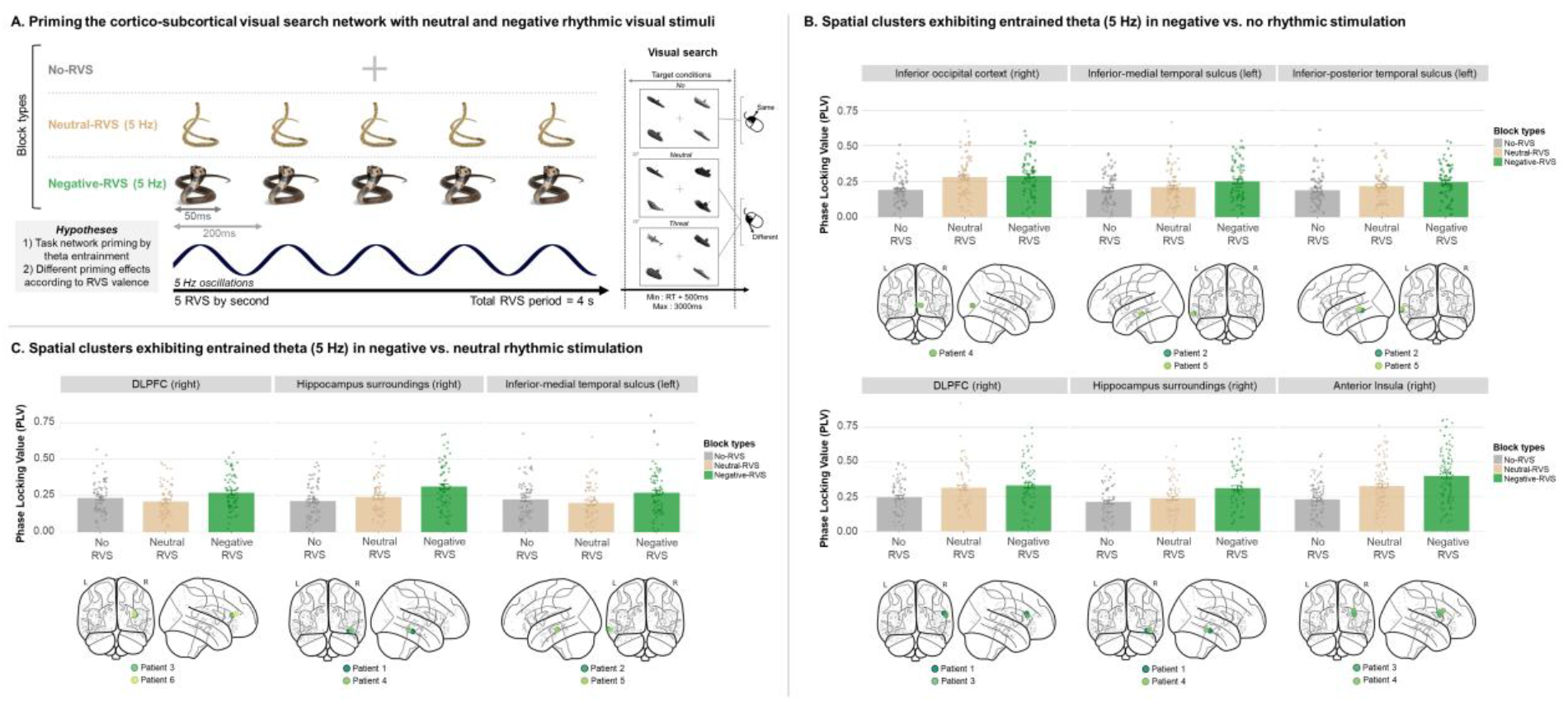
A) Blocks with no, neutral valence (rope), and negative valence (snake) 5 Hz RVS were used to prime the visual search network in a block design, presented before each trial. B) SEEG contacts (cluster-corrected) showing increased 5 Hz PLV in the negative valence-RVS compared to the No-RVS condition. C) SEEG contacts (cluster-corrected) showing increased 5 Hz PLV in the Negative-RVS compared to the Neutral-RVS condition. Bar plots display mean stim-brain PLV for each block type, with error bars indicating SEM and dots representing individual data points. Green dots on brain maps indicate the anatomical localization of significant clusters (paired t-tests).

Based on these results, we next questioned whether such stimulation can modulate behavioral performance in a group of healthy individuals.

### Negative theta RVS boosts attention and memory in healthy adults

We conducted a behavioral experiment in 27 healthy adults (mean age: 25.9 ± 3.4 years; S6) to investigate the impact of entrainment driven by theta Neutral-RVS (Rope/Control) and Negative-RVS (Snake/Threatening) on visual search performance. In this behavioral experiment, the visual search task was identical to that performed by SEEG patients but included a recognition block (episodic memory without stimulation) instead of a rest period at the end of the visual search blocks. Participants were not instructed to memorize items during the search task to test involuntary episodic learning and memory (Fig. 1 and 4.A). The recognition block (72 trials) included visual search targets (50%) and distractors (50%). Instructions for the recognition block were given only after the completion of the visual search task.

**Figure 4.**
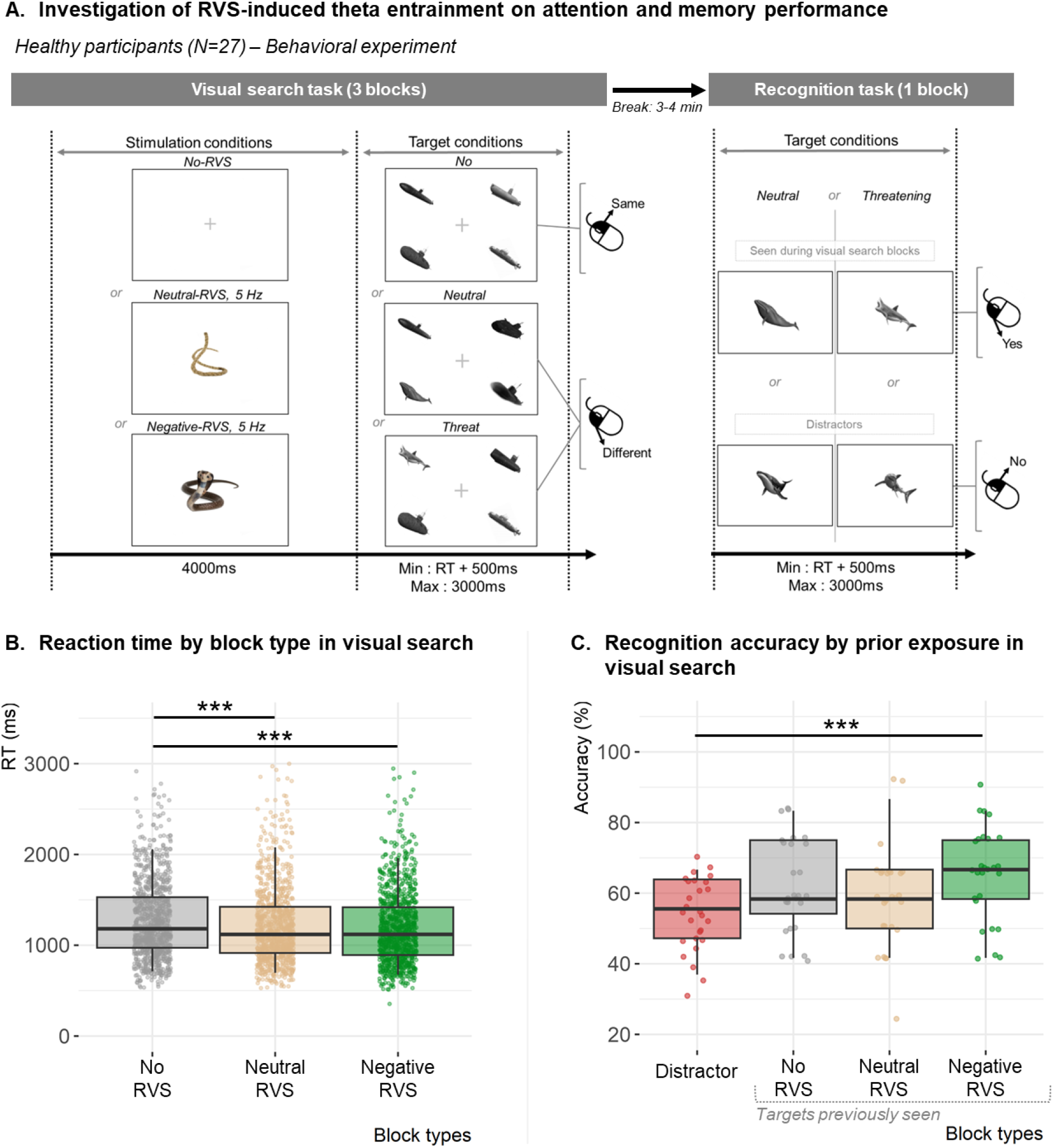
A) In the recognition task (right panel), participants were asked to determine whether a gray-shaded item had previously been presented as a target during the visual search blocks. Importantly, they were not informed in advance that their memory for these items would be tested. B) Median reaction time in the visual search task by stimulation conditions, and C) mean accuracy in the recognition task according to condition of prior exposure. In each boxplot, the horizontal line represents the median measure for the group, and the box shows the interquartile range. The vertical line indicates the 5th and 95th percentiles. Dots represent individual data. \*\*\**p* < .001.

In SEEG patients, Negative-RVS resonance propagated across networks involved in sensory and attention processing, as well as subcortical areas including the amygdala and the hippocampus, which play a notable role in fear learning and memory^19,25–28^. Thereby, we expected Negative-RVS to prime the task network, boosting both attention and memory. SEEG results of Neutral-RVS indicate a transitional phase-locking stage for RVS. We therefore hypothesized either a null or a minimal impact of Neutral-RVS on performance. As we were interested in investigating possible interaction between Neutral- and Negative-RVS and visual information to be detected, the target factor (No-, Neutral- and Threatening-Target) was considered in the following analysis.

Visual search accuracy averaged 91.4 ± 0.8%, and median RT of 1228.3 ± 30.9 ms; these variables were unaffected by age, sex, or handedness (Supplemental Information). No effect of Target or RVS condition was observed on accuracy. Target condition significantly affected reaction time (χ^2^ (2) = 29.9, *p* < .001): participants responded faster to Neutral- and Threat- Target trials than No-Target trials (both *p* < .001), with a superiority effect for Threat- over Neutral-Targets (*p* < .001). This effect was expected in healthy adults^29,30^, while disregarded in SEEG analysis because of limited sample size. Additionally, RT were faster in blocks with Neutral- and Negative-RVS compared to No-RVS (both *p* < .001, Fig. 4.B; RT Neutral-RV vs. Negative-RVS: *p* = .276, suggesting that theta RVS speeds up processing, irrespective of accuracy. This unspecific effect of RVS might optimize further neural framing of visual information into “perceptual moments” through theta oscillations^31,32^. This principle, also known as “theta phase precession”, is crucial for the hippocampus to encode spatial features, which in turn, enable efficient attention orienting during visual search^33^. In parallel, theta suppression in the insula has been associated with reduced arousal (physiological alertness) elicited by threat perception^14^. As we have observed in the SEEG data that 5 Hz Neutral- and Negative- RVS entrained theta in the insula (moderately and significantly, respectively, Fig. 3.B), we propose that 5 Hz RVS may prevent theta suppression by heightening arousal. This hypothesis aligns with behavioral effects observed in healthy adults, with faster responses for trials with RVS irrespective of valence. Importantly, the intermediate behavioral and neurophysiological effects observed under Neutral-RVS suggest that valence modulates entrainment in a graded rather than binary fashion. The absence of significant memory facilitation in the Neutral-RVS condition, despite modest theta entrainment, implies that stronger emotional salience may be required to engage subcortical circuits essential for episodic encoding.

Given the facilitating role of theta phase precession in encoding^31,32^, we next sought to investigate whether the entrainment induced by theta RVS, particularly in the Negative-RVS condition, could facilitate memory encoding of target stimuli during visual search, consequently enhancing participants’ episodic memory performance in the recognition task (Fig. 4.A). In the recognition task, the mean accuracy was 58.1 ± 0.9 % (min: 51.4 %, max: 70.8 %) and the mean median RT 1207.7 ± 36.6 ms. There was no effect of Target on RT (*p* = .068). However, stimulation had a significant effect on accuracy (χ ^2^ (3) = 15.8, *p* = .001) driven by increased recognition rates of items (threat or neutral) seen in the Negative-RVS block compared to distractors (*p* < .001, Fig. 4.C; all other contrasts were non-significant, with *p* = .062 for Distractor vs. No-RVS block as closest tendency). Based on the SEEG results, we propose that this facilitation effect might ensue from the theta phase precession phenomenon, promoting memory encoding during visual search, as suggested by higher accuracy for items seen in the Negative-RVS block compared to distractors.

## Discussion

In the present study, we combined SEEG recordings in epileptic patients with a behavioral experiment in healthy adults to determine whether 5 Hz negative-valence rhythmic visual stimulation could i) entrain cortical and subcortical structures (SEEG experiment) and ii) modulate attention and memory (behavioral experiment in healthy participants). Based on evidence that negative emotional content robustly engages limbic circuits^13–15,27^, we hypothesized that negative-valence stimulation would result in i) more pronounced theta entrainment than neutral stimulation in SEEG data, ii) stronger behavioral facilitation, than neutral stimulation and greater memory encoding in healthy individuals.

When no RVS preceded the visual-search period, we observed transient increases in both theta and gamma magnitude in occipital SEEG contacts, consistent with feature binding before visual information is routed along the dorsal (“where”) and ventral (“what”) pathways, respectively engaging the superior occipital and inferior temporal cortices^34–36^. This finding aligns with previous studies suggesting that theta and gamma oscillations co-occur in the visual cortex to support both early visual information processing^37,38^, with theta oscillations in the superior-posterior temporal gyrus notably supporting attentional orienting^39,40^.

We then showed that, in absence of stimulation, the spontaneous theta activity during visual search contained two distinct, but complementary, components: an initial transient burst (0-0.3 s post-visual search onset) in occipital and superior-posterior temporal cortices as well as in the amygdala, and a slow/sustained increase (0-1.5 s) in limbic regions (hippocampus, amygdala, anterior insula), orbitofrontal cortex, and prefrontal regions (dorsolateral and dorsomedial). The early transient theta responses in occipital and posterior temporal clusters is consistent with rapid visual feature binding and initial attentional orienting processes occurring in sensory cortices and adjacent temporal areas^34–38^. The simultaneous transient amygdala theta response might be related rapid emotional appraisal or salience detection mechanisms that guide immediate attentional prioritization^25,26,41^. In contrast, we propose that the slow and sustained theta increase in limbic and prefrontal regions is associated with ongoing associative and experience-dependent processing, reflecting a shift toward more integrative cognitive and emotional functions such as memory retrieval, decision-making, and executive control^26,27,40–42^.

Specifically, the gradual theta rise observed in the hippocampus highlights its roles in memory and spatial contextualization, particularly in binding items to their spatial and temporal context during the visual search^25,26,41^. We propose that the sustained theta responses observed in the anterior insula, orbitofrontal cortex, and particularly the dorsolateral prefrontal cortex, support the maintenance of attentional states^40^ and emotional evaluation^43^, as well as the accumulation and integration of perceptual evidence during visual search^36–38,44^. The temporal distinction between transient/sensory-driven theta responses and sustained activity in limbic and higher-order cortical areas provide novel insights into how visual search engages brain regions sequentially, transitioning from immediate perceptual analyses to more complex cognitive and emotional integration.

Based on these results observed without stimulation, and considering the well-established role of theta oscillations in attention and working memory^2,17–19,39,45,46^, we investigated whether these task-related oscillatory markers in the theta range could be entrained and modulated by rhythmic sensory stimulation. Participants were exposed to brief (4 s) blocks of flickering images, either a snake (Negative-RVS) or a rope (Neutral-RVS), presented at 5 Hz before each visual search trial. By manipulating emotional valence, we aimed to determine whether negative-valence rhythmic input would more effectively entrain cortico-subcortical networks critical for attentional and memory processes in the visual search task.

Negative-RVS entrained (Stim-Brain PLV during stimulation) theta across cortico-subcortical networks, notably the hippocampus and the dorsolateral prefrontal cortex, beyond what was observed for neutral stimuli. This valence-specific boost in theta entrainment suggests that negative RVS optimally synchronizes emotion and memory circuits, reinforcing phase alignment in limbic and prefrontal areas crucial for fear processing and episodic encoding. Our data thus reveal how negative-valence RVS is likely to tap into pre-existing emotion-memory pathways, amplifying theta synchrony between ventral visual and limbic regions that are primed for attentional detection. Together, the No-RVS, Neutral-RVS, and Negative-RVS conditions reveal a parametric gradient of neural entrainment, with valence serving as a functional modulator of theta synchronization strength. This graded profile reinforces the idea that valence dynamically tunes the depth of sensory-driven recruitment across cortico-subcortical networks.

Critically, the stronger entrainment induced by negative-valence stimulation compared to neutral stimulation supports the idea that emotional valence amplifies synchronization in subcortical networks, surpassing the more moderate effects observed with neutral rhythmic stimulation. Negative rhythmic stimulation thus appears to non-additively prime theta activity in the hippocampus and DLPFC, enhancing the interplay between top-down attention (DLPFC) and memory (hippocampus). We propose that this mechanism can possibly boost the accumulation of visual evidence prior to the motor response, and can potentially facilitate memory encoding in the ventral visual pathway^44–47^. To test this hypothesis, we performed a behavioral experiment in healthy individuals who performed the same task as SEEG patients with or without stimulation followed by a recognition task.

Behavioral results were well aligned with our SEEG-based hypotheses. We observed that both negative and neutral rhythmic stimulation speeded reaction times, presumably by helping structure perceptual moments through theta oscillations^33^, whereas negative valence specifically led to enhanced memory recognition. These results point to a mechanistic pathway in which threatening visual flickers preferentially synchronize limbic circuits at theta frequencies, potentially creating an ideal temporal window for enhanced encoding of salient or fear-relevant stimuli. Together with the progressively rising theta in hippocampus and DLPFC we reported in SEEG data (Fig. 2), these findings suggest that Negative-RVS accelerates the accumulation of visual evidence before response execution^48^ by sharpening ventral-pathway recognition processes^44,47^ while enhancing DLPFC-hippocampus dialogue that supports episodic encoding^44–47^.

Finally, the heightened memory for target items presented under negative rhythmic stimulation also converges with the principle of theta phase precession, which has been linked to stronger memory encoding in the hippocampus^31,32^. We propose that the engagement of emotional circuitry by negative-valence rhythmic stimulation capitalizes on the amygdala–hippocampal relationship implicated in learning and memory^19,25–28^, reinforcing the salience of relevant stimuli and enhancing subsequent recognition. Clinically, such selective memory benefits raise the prospect that valence-based rhythmic stimulation could serve as a non-invasive adjunct in disorders where memory or emotion regulation is compromised (e.g., post traumatic stress disorder^15^, anxiety^49^, or age-related cognitive decline^50,51^).

## Conclusion

By uniting rare intracranial SEEG recordings from epilepsy patients with a complementary behavioral experiment in healthy adults, we demonstrate that the valence of the rhythmic sensory stimulus is a key feature to powerfully entrain theta-band oscillations in both cortical and subcortical networks. Specifically, we report robust synchronization between negative valence stimulation and entrainment in the ventral visual stream, hippocampus, and dorsolateral prefrontal cortex, brain regions essential to sustaining attention and episodic encoding. Crucially, while both neutral and negative stimulation enhanced visual search performance, only negative-valence stimulation boosted recognition memory for subsequently encountered items.

To conclude, our findings carry broad translational significance: by highlighting the influence of emotional valence, future interventions may capitalize on rhythmic entrainment to refine attention and memory processes. This approach could indeed serve as a non-invasive complement to non-invasive electrical or magnetic stimulation techniques, offering new possibilities for engaging deep cortical and subcortical networks in clinical interventions. Ultimately, this work establishes valence not merely as an affective feature, but as a functional parameter of stimulation, capable of deepening neural entrainment and extending its reach into subcortical and associative cortical regions.

## Materials and Methods

### Resource availability

#### Lead contact

Further information and data that support the findings of this study are available upon request from the corresponding author, Roxane S. Hoyer.

#### Material availability

The stimuli and codes used to build and present the behavioral task are available online on OSF (https://osf.io/g4ap5/). The raw brain data used in this study (MRI and SEEG recordings) are protected as acquired through clinical assessment and are not available for sharing (2022- 5890). Pre-processed SEEG data used to generate the figures and MATLAB scripts are available at the following URL(https://osf.io/g4ap5/).

#### Data and code availability

Original codes used to process SEEG and behavioral data are publicly available on OSF (https://osf.io/g4ap5/), or upon request to the corresponding author.

### Experimental design and populations

#### Patients

SEEG data from 6 patients affected by drug-resistant focal epilepsy were collected (mean age and standard deviation: 30 ± 3.9 years; male: 50 %, female: 50 %) following the Research Opportunities in Humans Consortium guidelines^52^. All patients performed the task above chance. The study protocol was described verbally by the research and clinical teams and the inclusion was discussed prior to hospitalization with patients. All patients provided verbal assent and written informed consent. This research complies with all relevant ethical regulations; the experimental procedures were approved by the Ethics Review Board of the CHU de Québec - Université Laval (2022-5890, Québec, QC, Canada).

#### Healthy participants

Behavioral data from 27 neurologically healthy young adults were collected and analyzed (mean age: 25.9 ± 3.4 years; male: 52 %, female: 48 %). All participants reported normal or corrected vision and hearing, in addition to no history of neurological or psychiatric disorder. They gave written informed consent and received a monetary compensation for their participation. All participants provided verbal assent and written informed consent. This project was approved by the Ethics Review Board of the CIUSSS de la Capitale Nationale (2022- 2476, Québec, QC, Canada).

### Method details

#### Behavioral tasks

##### SEEG experiment

In visual search trials, patients viewed four items in grey shades and had to quickly determine if all items were from the same category (e.g., 4 submarines) or if one item was different (e.g., 3 submarines and 1 whale) by pressing the right or left mouse key. They had 3 s to respond before the next trial. All task blocks included equal proportions of No-Target (e.g., 4 submarines), Neutral-Target (e.g., 3 submarines and 1 whale), and Threatening-Target (e.g., 3 submarines and 1 shark) trials. Both living and non-living items were used as targets (Fig. 1.A).

In the No-RVS block, each trial was preceded by a 4 s fixation cross. In the Neutral-RVS block and Negative-RVS block, trials were preceded by a 4 s display of 5 Hz flickering picture of a rope or a snake (50 ms duration, 200 ms inter-onset interval), respectively (see Fig. 3). The negative and neutral stimuli were projected on a screen (1024*768 pixels located 50cm in front of the participant). The duration of each block (36 trials) varied as a function of participants’ response times (minimum block duration: 3 min 36 s, maximum: 5 min 24 s). The task was implemented using a block design, with blocks randomization across patients. Target types and positions were balanced within and across task blocks. Once the task performed, patients’ brain activity was recorded during an open-eyes resting state period (5 min 24 s). Overall, brain activity in SEEG patients was recorded across four blocks: No-RVS, Neutral valence-RVS, Negative valence-RVS, and Rest.

Patients were informed they were going to perform an attention task, including two blocks with trials preceded by flickering images before each trial. They were instructed to passively observe pictures (Rope-Neutral valence and Snake-Negative valence RVS) flickering at 5Hz before each task trial; if none were presented (No-RVS condition), they were asked to keep fixating the center of a blank screen. Participants were instructed to indicate as quickly as possible if all items belonged to the same category (right mouse button for “yes,” left for “no”).

##### Healthy participants

Healthy adults performed the exact same task and blocks as SEEG patients, but the rest period was replaced by a memory-recognition block, where they were shown targets that had been presented in the visual search blocks (Neutral- and Threatening- Targets) alongside distractors (Fig. 3.A). Each trial of the recognition task began with a 2 s fixation cross, followed by a single item which participants had 3 s to determine if this item has been previously seen or not seen during the visual search task. In 50% of the trials, the grey-shaded item, was a visual search target. We used an equal number of neutral and threatening, and living and non-living, stimuli. In the other 50%, the item was a distractor, slightly different from the visual search task targets. In the recognition block, the distractors were matched to targets’ valence (neutral, threatening) and nature (living, non-living). This block was challenging, as participants were not informed beforehand that they will perform a recognition task using the targets from the visual search blocks (total of 36 target and 36 distractor trials). Participants were therefore encouraged to perform accurately but to give intuitive responses if unsure.

#### Procedure

SEEG patients, with electrodes implanted based on clinical requirements, were tested at the Hôpital de l’Enfant-Jésus, CHU de Québec (QC, Canada) in a dedicated SEEG Epilepsy Monitoring Unit. SEEG data were recorded continuously, and patients were seated approximately 50 cm from the computer screen. Prior to the recording, each patient received verbal and illustrated instructions to ensure task comprehension, without practice trials. Presentation software (Neurobehavioral Systems, Albany, CA, USA) was used to present visual stimuli and record behavioral responses. They completed three VS blocks of 36 pseudo-randomized trials each; block order was randomized across Stimulation conditions (No-RVS, Neutral-RVS, Negative-RVS). Patients were instructed to maintain focus on the screen and respond as fast as possible using the mouse keys. After completing the VS task blocks, patients underwent a resting-state recording period.

Healthy participants were tested individually in a quiet room with the same procedure and apparatus. During the task, they seated in front of a computer (∼50 cm from the screen) in a quiet room. Before performing the VS task (same procedure), participants received detailed verbal and illustrated instructions (no training). Following these blocks, healthy participants performed the recognition task immediately after the VS task.

#### SEEG electrodes and contacts localization

Patients’ brains were stereotactically implanted with 16 to 18 semi-rigid multi-lead electrodes (AdTech, diameter: 0.8 mm). According to the clinically defined targeted structures, brain activity was recorded over SEEG contacts from of 2.0 mm wide and 3, 4, 5 or 6 mm apart (depending on the electrode). A 3D anatomical MPRAGE T1-weighted MRI (3T Siemens Trio, Siemens AG, Erlangen, Germany) was recorded pre-implantation of SEEG electrodes, with 160 sagittal slices and voxels of 1mm^3^ covering the whole brain. Cortical surfaces were extracted from the T1-weighted anatomical MRI and electrodes’ contacts localization was defined using the post-implantation CT scan co-registered with the pre-implantation MRI (ImaGIN toolbox, https://f-tract.eu/software/imagin/). MNI coordinates were computed using the SPM toolbox (http://www.fil.ion.ucl.ac.uk/spm/) and the SEEG contacts localization was defined based on the AAL3 atlas^53^.

#### SEEG recordings and preprocessing

Intracranial brain activity was simultaneously recorded from SEEG contacts (sampling rate: 512 Hz) using a video-SEEG monitoring system (Natus). Signals were initially referenced to a SEEG contact located in the white matter during acquisition. The signals were bandpass filtered from 0.1 to 200 Hz during acquisition. Contacts or trials containing interictal epileptiform discharges or high-amplitude noise were identified by visual inspection and rejected in collaboration with the clinical team. A notch filter was applied to reduce powerline contamination of the raw data (main 60 Hz, harmonics 120 and 160 Hz). To analyze brain activity before motor response to the task (mean median RT in patients: 1623 ± 112.3, minimum: 1278 and maximum: 2084), SEEG signals were epoched to consider trials extending from -4 s pre- to 2 s post-search onset (baseline: -2 to 0 s or -6 to -4s depending on further analysis to be performed). Each SEEG signal was then re-referenced to its immediate neighbor through bipolar derivations. Bipolar montage refines local specificity by eliminating signal artifacts common to adjacent contacts (e.g., 60 Hz running through wires and distant physiological artifacts) and cancel effects of distant sources whose signal is spreading to adjacent sites through volume conduction. Using this method, the estimated spatial resolution of each SEEG bipolar contact was of 3-4 mm^54^. Across patients, the brain activity was recorded over a total of 77 to 84 bipolar contacts (see SI S1).

#### SEEG signal processing and analyses

*Signal preprocessing*. SEEG preprocessing and statistical analyses were performed using Brainstorm (http://neuroimage.usc.edu/brainstorm/)^55^, FieldTrip functions (www.fieldtriptoolbox.org), and custom MATLAB codes.

##### Morlet wavelets

To evaluate the magnitude of task-related brain oscillatory activity during visual search and RVS, time-frequency Morlet decompositions (wavelet transform of the signal) were performed for each SEEG contact, each trial and each participant. The wavelet family was defined by (f0 /sf) = 3 with f0 ranging from 1 to 150 Hz with 1 Hz steps. Time frequency maps were normalized (z-score) according to the mean magnitude computed over the baseline period (−2 to 0 s for No-RVS vs. Rest analysis, -6 to -4s for No-RVS vs Neutral- RVS or Negative-RVS analysis). Two tailed Paired T-tests with Monte Carlo simulations (1000) and FDR correction for multiple testing over time and frequencies (alpha threshold: *p* < .05) were performed for each SEEG contact of each participant (during the response period in No- RVS vs. Rest and during the stimulation and response period in No-RVS vs Neutral-RVS or Negative-RVS).

##### Hilbert transform

To identify time related fluctuations of oscillatory rhythms associated with task engagement (No-RVS vs. Rest, see aforementioned analysis), we used the Hilbert transform to extract magnitude of theta oscillations (see results for rational). We first band- pass filtered the signal of each SEEG contact and each trial between 4 and 8 Hz (theta) and extracted its envelope. Bandpass filtering for theta was performed using a zero-phase 4th order Butterworth filter, with edge-trial padding of 1s to minimize edge artifacts. Hilbert data were normalized (z-score) over the baseline (−2 to 0 s for No-RVS vs. Rest). Significant differences in theta magnitude across task conditions were assessed using two-tail paired T-tests (averaged over time, 0 to 2 s, visual search period) with Monte Carlo simulations and FDR correction for multiple testing over time (alpha threshold: *p* < .01) were applied.

##### Spatial clustering

To identify the brain areas supporting visual search and Stim-Brain PLV the MNI coordinates of SEEG contacts showing significant effects were extracted. For all patients, significant contacts were projected over a standardized brain space (MNI), and a 3- dimensional sphere was projected on the MRI volume showing a significant increase in oscillatory magnitude (radius: 4 mm). The radius size was defined based on the estimated spatial resolution associated with bipolar montage. Only clusters of at least two overlapping SEEG contacts were considered as significant for all analyses, while isolated ones were disregarded (minimization of type I error rate). This analysis was performed using custom MATLAB codes (see “data and code availability” section). Spatial clusters were labelled based on their location according to the AAL3 altlas in the standard ICBM32 brain. Clusters of significant contacts in the anterior and medial middle temporal gyrus were respectively named “amygdala surroundings” and “hippocampus surroundings” due to their proximity with these subcortical areas (see SI S3).

##### Time series clustering

To identify task-related functional networks, we analyzed theta (4–8 Hz) magnitude (Hilbert) using a dynamic time warping (DTW) analysis, combined with the Partitioning Around Medoids (PAM) algorithm^56,57^. DTW computes the similarity between temporal sequences by aligning time series to minimize phase and amplitude differences, making it well-suited for capturing variations in oscillatory dynamics across regions. We used this method to group theta time series with similar temporal profiles, allowing us to identify dissociated transient and sustained patterns of activity, which may reflect the engagement of distinct functional networks (see results). Given that visual search involves both early, transient processes linked to attentional orienting and recognition, and sustained processes associated with higher-order integration and prediction^20–23^, the unsupervised algorithm automatically identified two clusters reflecting this expected biphasic structure in the theta dynamics. Temporal clustering was performed using the “kmedoids” MATLAB function^58^, which minimizes the sum of dissimilarities within clusters by assigning each time series to its closest medoid. This analysis was implemented using the aforementioned function and custom MATLAB codes (see “data and code availability” section).

##### Phase-Locking Value

We then computed the Phase Locking Value (PLV), between the patients’ 5 Hz oscillations and a theoretical pure 5 Hz sinusoid over the trial period. The phase-locking value (PLV) measures the magnitude of the average phase difference between two signals represented as a unit-length complex vector^24^. By presenting 5 Hz RVS before the target occurrence in two of the three task blocks, we aimed at entraining brain oscillatory activity at this frequency to prime the task network. To test this hypothesis, we performed a two-tail paired T-test between conditions (No-RVS, Neutral-RVS, Negative-RVS; positive direction with values averaged over time) with *p* < .05 as significance threshold. This analysis was performed using the Brainstorm MATLAB toolbox. The period considered ranged from 0 to 2 s following visual search onset. A baseline period from -6 to -4 s (pre-stimulation epoch) was used for normalization in comparisons between conditions (No-RVS, Neutral-RVS, and Negative-RVS).

#### Behavioral data analyses

SEEG patients correctly performed the task (mean accuracy ± SEM – No-RVS: 83.8 ± 2.5 %; Neutral-RVS: 86.1 ± 3.0 %; Negative-RVS: 84.3 ± 5.3 %) but their performance as a function of tragets and stimulation conditions was not analyzed due to the small sample size.

In healthy participants successfully performed the task (mean accuracy ± SEM – No-RVS: 91.8 ± 1.2 %; Neutral-RVS: 89.8 ± 1.2 %; Negative-RVS: 92.4 ± 1.6 %). RT (> 0 ms) from correct responses were log transformed to better fit a gaussian distribution (Shapiro-Wilk test: W = 0.91, *p* < .001). RTs were log-transformed to improve normality and analyzed using a linear mixed-effects model, with Participant as a random effect and Target, Block, Age, Sex, and Handedness as fixed effects. Accuracy was analyzed using a generalized linear model with binomial distribution. RT and accuracy to the visual search task were modeled using lmer (RT: gaussian fitting) and glm (accuracy: binomial fitting) R^59^ functions (lme4 and stats packages), respectively. Five fixed factors were considered in the analysis: Target (No, Neutral and Threatening), Block (No, Neutral- or Negative-RVS), Age (numeric vector), Biological Sex (Male and Female) and Handedness (Right or Left). Target and Block were modeled with possible interaction (variables of interest), while only the main effects of Age, Biological Sex and Handedness, were considered (between-participants variables with possible cofounding effects). Finally, Participant was used as random factor in RT analysis (lmer R function), but not in accuracy (glm R function), still to avoid overfitting issues leading to singular fit, low generalization power, and high variance in predictions. The same method was applied to analyze accuracy in the recognition task (Target levels: distractor, items seen in the Neutral- RVS block, and items seen in the Negative-RVS block). Type-II Wald tests (car package) were performed on the models. P-values were considered significant at *p* < .05. When a significant main effect or interaction was found, post-hoc Honest Significant Difference (HSD) pairwise tests were performed (emmeans package). In the case of multiple comparisons, the p-value of the Tukey HSD test was adjusted using the Tukey method for comparing multiple estimates (see also SI S6 for details on the methods used to analyze behavioral data).

## Acknowledgements

This work was supported by a Post doctoral fellowship from the Fonds de Recherche du Québec – Nature et Technologies to R.S.H, a grant from Fonds de Recherche du Québec - Santé and Brain Canada Future Leaders to P.A., a grant from Réseau de Bio-Imagerie du Québec to P.A., and NSERC Discovery grant to I.B. and P.A.

## Authors contribution

Conceptualization, R.S.H., I.B., A.Bl. and P.A.; Methodology, R.S.H, L.M., P.L-B., I.B., A.Bl. and P.A.; Software, R.S.H., A.Bo., L.C. and P.A.; Formal Analysis, R.S.H. and P.A.; Investigation, R.S.H., M-F.B., L.M., P.L-B., A.Bo.; Resources, L.M., P.L-B. and P.A.; Data Curation, R.S.H., M-F.B., and A.Bo.; Writing - Original Draft, R.S.H.; Writing - Review and Editing, R.S.H, M-F.B, L.M., P.L-B., C.L., A.Bo., I.B., A.Bl., S.L. and P.A.; Visualization, R.S.H., L.C.; Supervision, P.A.; Funding Acquisition, R.S.H., I.B., S.L. and P.A.

## Inclusion and diversity

We are committed to fostering inclusive, diverse, and equitable practices in all aspects of our research, ensuring that our findings contribute to a broader understanding of human neuroscience.

## Data and code availability

The material used in this study, including codes and behavioral data from healthy participants are available online (https://osf.io/g4ap5/), or upon request by contacting Roxane S. Hoyer. The raw brain data used in this study (MRI and SEEG recordings) are protected as acquired through clinical assessment and are not available for sharing (2022-5890); averaged data are available online.

