## Supplemental information for "Valence of sensory stimulation as a key feature for subcortical entrainment: Insights from human intracranial EEG"

### S1. Patients' characteristics

| | Biological sex | Handedness | Age in years | Education | Number of implanted electrodes | Total of bipolar contacts |
|---|---|---|---|---|---|---|
| ***Patient 1*** | Male | Right | 29 | Vocational diploma | 17 | 79 |
| ***Patient 2*** | Male | Right | 26 | Secondary 5 | 16 | 83 |
| ***Patient 3*** | Female | Right | 34 | Vocational diploma | 18 | 77 |
| ***Patient 4*** | Female | Right | 25 | Secondary 5 + 1 vocational diplomas | 18 | 77 |
| ***Patient 5*** | Female | Right | 32 | Secondary 4 + 2 vocational diplomas | 18 | 80 |
| ***Patient 6*** | Male | Right | 34 | Secondary 4 + vocational diploma | 18 | 84 |

### S2. SEEG patients' performance to the visual search task

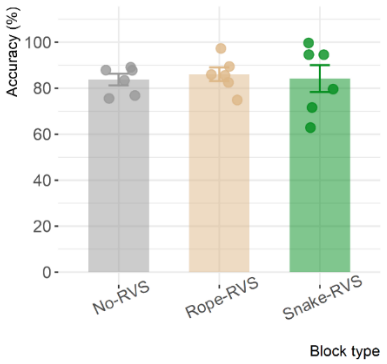


Figure S2.1. Accuracy by stimulation condition in SEEG patients. The barplots represent the mean accuracy (%) across three stimulation conditions: No-RVS (gray), Neutral-RVS (beige), and Negative-RVS (green). Error bars indicate the standard error of the mean (SEM). Each point represents the accuracy for an individual patient (N = 6).

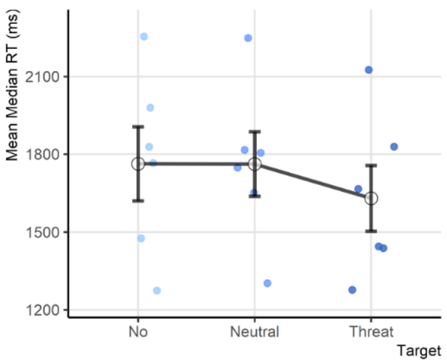


Figure S2.2. Mean median reaction Time (RT) by target conditions in No-RVS Trials. Target conditions: "No" (light blue), "Neutral" (medium blue), and "Threat" (dark blue) during No-RVS trials. Each black dot represents the mean median RT (ms) across participants, with error bars indicating the standard error of the mean (SEM). Individual participant data points are displayed as blue shaded jittered dots. The black line connects the mean RT values across conditions to highlight trends.

### S3. Theta network supporting visual search: t-test Hilbert (FDR *p* < .01)

Table S3. Significant SEEG contacts for the No-RVS vs. Rest comparison for Hilbert theta power analysis, with anatomical localization and related label. The first table lists significant contacts across spatial clusters, including electrode names, corresponding anatomical labels (AAL3), MNI coordinates for each patient, and the number of significant contacts across clusters for each patient.

| Spatial cluster | Regroupment | Patient | Electrode name | AAL3 | MNI coordinates | | |
|---|---|---|---|---|---|---|---|
| | | | | | *x* | *y* | *z* |
| DMPFC (right) | | Patient 1 | Z1-Z2 | Vent_Str_R | 4 | 52 | 13 |
| | | Patient 1 | Z3-Z4 | Frontal_Sup_Medial_R | 7 | 53 | 20 |
| | | Patient 1 | Z5-Z6 | Frontal_Sup_Medial_R | 10 | 53 | 26 |
| | | Patient 1 | Z7-Z8 | Frontal_Sup_2_R | 13 | 51 | 31 |
| DPLFC (right) | | Patient 1 | Q5-Q6 | Frontal_Inf_Oper_R | 45 | 18 | 15 |
| | | Patient 3 | Q3-Q4 | Frontal_Inf_Tri_R | 43 | 25 | 9 |
| | | Patient 3 | G6-G7 | Frontal_Inf_Tri_R | 37 | 31 | 12 |
| | | Patient 3 | Q1-Q2 | Frontal_Inf_Tri_R | 36 | 24 | 7 |
| | | Patient 3 | G8-G9 | Frontal_Inf_Tri_R | 51 | 29 | 9 |
| | | Patient 3 | G9-G10 | Frontal_Inf_Tri_R | 58 | 28 | 9 |
| | | Patient 3 | Q7-Q8 | Frontal_Inf_Tri_R | 55 | 25 | 14 |
| | | Patient 5 | H8-H9 | Frontal_Inf_Tri_R | 50 | 26 | 7 |
| Insula anterior (left) | | Patient 2 | Xp3-Xp4 | Insula_L | -33 | 15 | -4 |
| | | Patient 3 | Xp3-Xp4 | Insula_L | -33 | 14 | -6 |
| | | Patient 5 | Xp1-Xp2 | Insula_L | -37 | 12 | -12 |
| | | Patient 5 | Xp3-Xp4 | Insula_L | -35 | 13 | -6 |
| Occipital inferior (right) | | Patient 4 | OI1-OI2 | Occipital_Sup_R | 3 | -79 | 3 |
| | | Patient 4 | OI3-OI4 | Cuneus_R | 11 | -78 | 3 |
| | | Patient 4 | OS3-OS4 | Lingual_R | 12 | -72 | 21 |
| | | Patient 4 | OS5-OS6 | Lingual_R | 22 | -74 | 25 |
| | | Patient 4 | OS7-OS8 | Occipital_Mid_R | 27 | -70 | 31 |
| OFC (left) | | Patient 2 | Op3-Op4 | Rectus_L | -15 | 34 | -15 |
| | | Patient 5 | Gp3-Gp4 | White_Matter | -19 | 34 | -12 |
| OFC (right) | | Patient 3 | O3-O4 | Rectus_R | 10 | 31 | -16 |
| | | Patient 3 | O5-O6 | White_Matter | 16 | 32 | -13 |
| | | Patient 3 | O7-O8 | OFCant_R | 22 | 32 | -11 |
| | | Patient 5 | O1-O2 | White_Matter | 3 | 30 | -28 |
| | | Patient 6 | G1-G2 | ACC_sup_R | 8 | 30 | -9 |
| | | Patient 6 | O1-O2 | Rectus_R | 7 | 33 | -20 |
| Temporal inferior - posterior (left) | | Patient 5 | Ap5-Ap6 | Cerebelum_Crus2_L | -45 | -6 | -29 |
| | | Patient 5 | ETp7-ETp8 | Cerebelum_Crus2_L | -52 | -7 | -33 |
| Temporal superior - posterior (right) | | Patient 4 | U3-U4 | Temporal_Mid_R | 44 | -27 | 5 |
| | | Patient 4 | U5-U6 | Temporal_Mid_R | 51 | -25 | 6 |
| Amygdala (right) | | Patient 2 | B3-B4 | ParaHippocampal_R | 32 | -11 | -24 |
| | | Patient 5 | B3-B4 | Amygdala_R | 32 | -15 | -24 |
| Middle temporal gyrus - anterior (left) | Amygdala surroundings (left) | Patient 1 | Bp5-Bp6 | Cerebelum_Crus2_L | -45 | -13 | -22 |
| | | Patient 1 | Bp7-Bp8 | Temporal_Inf_L | -53 | -13 | -23 |
| | | Patient 2 | Tp3-Tp4 | Temporal_Mid_L | -50 | -2 | -11 |
| | | Patient 2 | Tp6-Tp7 | Temporal_Mid_L | -60 | 2 | -10 |
| | | Patient 5 | Tp1-Tp2 | White_Matter | -45 | -11 | -17 |
| | | Patient 5 | Tp3-Tp4 | Temporal_Inf_L | -51 | -9 | -15 |
| | | Patient 5 | Tp5-Tp6 | Temporal_Inf_L | -57 | -7 | -12 |
| | | Patient 5 | Tp7-Tp8 | Temporal_Inf_L | -61 | -6 | -11 |
| | | Patient 3 | Ap5-Ap6 | White_Matter | -38 | -1 | -21 |
| | | Patient 3 | Ap7-Ap8 | White_Matter | -45 | 0 | -21 |
| | | Patient 3 | Ap9-Ap10 | Temporal_Inf_L | -53 | 1 | -21 |
| Middle temporal gyrus - anterior (right) | Amygdala surroundings (right) | Patient 1 | A7-A8 | Temporal_Inf_R | 51 | -3 | -27 |
| | | Patient 1 | A8-A9 | Temporal_Inf_R | 57 | -2 | -26 |
| Middle temporal gyrus - medial (left) | Hippocampus surroundings (left) | Patient 2 | Cp9-Cp10 | Temporal_Inf_L | -54 | -27 | -15 |
| | | Patient 5 | Bp5-Bp6 | Temporal_Inf_L | -45 | -22 | -15 |
| Middle temporal gyrus - medial (right) | Hippocampus surroundings (right) | Patient 1 | C3-C4 | White_Matter | 41 | -28 | -16 |
| | | Patient 1 | C5-C6 | White_Matter | 48 | -28 | -13 |
| | | Patient 1 | C7-C8 | Temporal_Inf_R | 56 | -28 | -12 |
| | | Patient 4 | B7-B8 | Temporal_Inf_R | 55 | -36 | -8 |

| Patient | Patient 1 | Patient 2 | Patient 3 | Patient 4 | Patient 5 | Patient 6 |
|---|---|---|---|---|---|---|
| **Number of significant contacts across clusters** | 12 | 6 | 13 | 8 | 13 | 2 |

### S4. Theta time course during visual search in significant spatial clusters

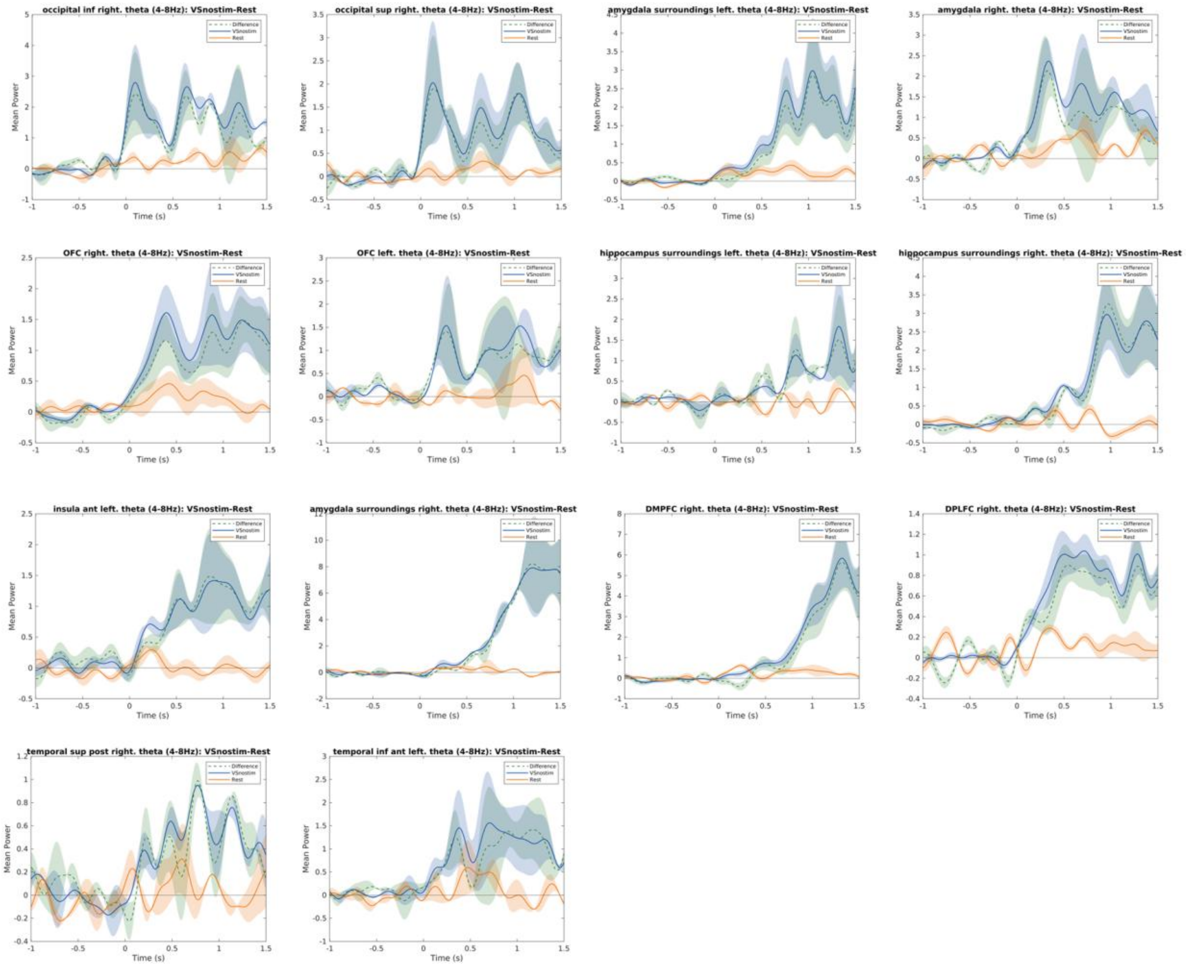


Figure S4. Theta time course (4-8 Hz) during visual search by spatial clusters. Each subplot represents theta power changes over time in a specific region of interest identified as significant through spatial cluster analysis. Data are shown for two conditions: visual search without stimulation (No-RVS, blue line) and Rest (orange line); the difference between values for these conditions is depicted by the dotted line (green line). Shaded regions indicate the standard error of the mean (SEM). Time zero marks the onset of the visual search task, with negative values representing pre-stimulus baseline. The clusters displayed correspond to significant differences in theta power (t-test, FDR corrected, all $p < .01$) between conditions during the task.

### S5. Theta (5 Hz) priming for visual search: t-test PLV (FDR *p* < .05)

#### *Negative-RVS vs. No-RVS*

Table S5.1. Significant SEEG contacts for the Negative-RVS vs. No-RVS comparison for PLV analysis, with anatomical localization and related label. The first table lists significant contacts across spatial clusters, including electrode names, corresponding anatomical labels (AAL3), MNI coordinates for each patient, and the number of significant contacts across clusters for each patient.

| Spatial cluster | Patient | Electrode name | AAL3 | MNI coodinates | | |
|---|---|---|---|---|---|---|
| | | | | *x* | *y* | *z* |
| DLPFC (right) | Patient 1 | Q 9-Q 10 | Frontal_Inf_Tri_R | 55 | 24 | 20 |
| | Patient 3 | Q 8-Q 9 | Frontal_Inf_Tri_R | 58 | 25 | 15 |
| Hippocampus surroundings (right) | Patient 1 | C3-C4 | White_Matter | 41 | -28 | -16 |
| | Patient 4 | B5-B6 | White_Matter | 45 | -35 | -14 |
| Insula anterior (right) | Patient 3 | X7-X8 | White_Matter | 34 | 9 | 15 |
| | Patient 3 | X9-X10 | White_Matter | 32 | 9 | 21 |
| | Patient 4 | H5-H6 | Frontal_Inf_Tri_R | 32 | 16 | 24 |
| Occipital inferior (right) | Patient 4 | OI1-OI2 | Occipital_Sup_R | 3 | -79 | 3 |
| | Patient 4 | OI3-OI4 | Cuneus_R | 11 | -78 | 3 |
| Inferior-medial temporal sulcus (left) | Patient 2 | Cp11-Cp12 | Temporal_Inf_L | -63 | -26 | -13 |
| | Patient 5 | Bp9-Bp10 | Temporal_Inf_L | -65 | -25 | -14 |
| Inferior-posterior temporal sulcus (left) | Patient 2 | Fp11-Fp12 | Temporal_Inf_L | -67 | -45 | -6 |
| | Patient 5 | Cp9-Cp10 | Temporal_Inf_L | -66 | -40 | -3 |

| Patient | Patient 1 | Patient 2 | Patient 3 | Patient 4 | Patient 5 | Patient 6 |
|---|---|---|---|---|---|---|
| **Number of significant contacts across clusters** | 2 | 2 | 3 | 4 | 2 | 0 |

#### *Negative-RVS vs. Neutral-RVS*

Table S5.2. Significant SEEG contacts for the Negative-RVS vs. No-RVS comparison for PLV analysis, with anatomical localization and related label. The first table lists significant contacts across spatial clusters, including electrode names, corresponding anatomical labels (AAL3), MNI coordinates for each patient, and the number of significant contacts across clusters for each patient.

| Spatial cluster | Patient | Electrode name | AAL3 | MNI coodinates | | |
|---|---|---|---|---|---|---|
| | | | | *x* | *y* | *z* |
| DLPFC (right) | Patient 3 | G6-G7 | Frontal_Inf_Tri_R | 37 | 31 | 12 |
| | Patient 6 | H6-H7 | Frontal_Mid_2_R | 33 | 37 | 14 |
| Hippocampus surroundings (right) | Patient 1 | C3-C4 | White_Matter | 41 | -28 | -16 |
| | Patient 4 | B5-B6 | White_Matter | 45 | -35 | -14 |
| Inferior-medial temporal sulcus (left) | Patient 2 | Cp13-Cp14 | White_Matter | -69 | -26 | -13 |
| | Patient 5 | Bp9-Bp10 | Temporal_Inf_L | -65 | -25 | -14 |

| Patient | Patient 1 | Patient 2 | Patient 3 | Patient 4 | Patient 5 | Patient 6 |
|---|---|---|---|---|---|---|
| **Number of significant contacts across clusters** | 1 | 1 | 1 | 1 | 1 | 1 |

### S6. Details on models used to analyze behavioral data

#### *Models' comparison*

To model the relationship between reaction times (RT) and the factors considered in this study, two linear mixed-effects models were compared. Model 1 used log-transformed RT as the dependent variable to address the non-normality of the raw RT data ($p < 0.001$, Shapiro-Wilk test), while Model 2 used the raw RTs. The fixed effects in both models included Block type (reference: No-RVS), Target (reference: No-Target), Sex (reference: Female), Handedness (reference: Left), and Age (a numeric variable). A random intercept for each participant was included to account for individual variability. The models were fitted using the lmer() function from the lme4 package in R. Model comparison was performed using the Bayesian Information Criterion (BIC), as the goal was to identify a parsimonious model that balances model fit with complexity (high penalization for inclusion of unnecessary parameters).

Model comparison using the BIC showed a strong preference for the model using log-transformed reaction times (Model 1). Model 1 had a BIC of 1274, significantly lower than Model 2, which used raw RTs (BIC = 42789). Given that BIC prioritizes parsimony and penalizes model complexity more strongly, these results support the choice of using log-transformed RTs to address the non-normal distribution of reaction times. All subsequent analyses were based on Model 1.

#### *Reaction times (RT) modeling*

*Visual search.* To analyze RT in healthy participants, we fitted a linear mixed-effects model using the log-transformed RT in ms as the dependent variable. The fixed effects included Block type and Target with interaction; Sex, Handedness, and Age, were included in the model in an exploratory approach, regarding to their possible main effect on data. A random intercept for each participant was considered. The marginal $R^2$ was 0.23, indicating that 23% of variance was explained by fixed effects, and conditional $R^2$ was 0.07. The residual variance was 0.09.

Table S6.1. Detail from models used to analyze RT. The table presents the fixed effects of the model, including estimates (β), standard errors (SE), degrees of freedom, t-values, and p-values. P-values indicate significance levels, with "< .001", "< .01", and "n.s." denoting not significant results.

| Variables | Estimate | Standard error | Degree of freedom | T-value | P-value |
|---|---|---|---|---|---|
| *Intercept* | 7,305 | 0,237 | 23,203 | 30,790 | < .001 |
| *Block type: Rope-RVS* | -0,071 | 0,023 | 2860,033 | -3,059 | < .01 |
| *Block type: Snake-RVS* | -0,060 | 0,023 | 2860,056 | -2,594 | < .01 |
| *Target: Neutral* | -0,118 | 0,023 | 2860,031 | -5,093 | < .001 |
| *Target: Threatening* | -0,197 | 0,023 | 2860,006 | -8,534 | < .001 |
| *Sex: Male* | -0,001 | 0,054 | 22,991 | -0,017 | n.s. |
| *Handedness: Right* | -0,039 | 0,077 | 23,008 | -0,509 | n.s. |
| *Age* | -0,003 | 0,008 | 23,000 | -0,322 | n.s. |
| *Block type: Rope-RVS*Target: Neutral* | 0,035 | 0,033 | 2860,011 | 1,081 | n.s. |
| *Block type: Snake-RVS*Target: Neutral* | -0,021 | 0,033 | 2860,027 | -0,642 | n.s. |
| *Block type: Rope-RVS*Target: Threatening* | 0,026 | 0,033 | 2860,002 | 0,790 | n.s. |
| *Block type: Snake-RVS*Target: Threatening* | -0,011 | 0,033 | 2860,012 | -0,337 | n.s. |

*Recognition*. RT were analyzed using the method described for the visual search blocks. No significant results were found.

*Accuracy modeling*

*Visual search*. To analyze accuracy during visual search (binary outcome: 0 = incorrect, 1 = correct), a generalized linear mixed-effects model (GLMM) was used with a binomial distribution and a logit link function. The model included Block_type (with "No-RVS" as the reference category) and Target (with "No-Target" as the reference category), as well as their interaction (Block_type*Target), as fixed effects to examine predictors of interest for the study. A random intercept for each participant was used. Given the binary nature of the dependent variable and the risk of overfitting due to the limited number of predictors relative to the dataset size, only the main predictors of interest were included in the model. The model was fit using the glmer function from the lme4 package in R. The bobyqa optimizer was employed with an increased maximum iteration limit (maxfun = 100,000) to ensure convergence. This approach ensured the model was stable.

*Recognition*. We used a generalized linear model (GLM) with a binomial distribution and logit link function to analyze accuracy in the recognition block. The fixed effects included the prior exposure factors (reference: “distractor”), Age (continuous), Handedness (reference: “Left”), and Sex (reference: “Female”). Initially, a generalized linear mixed-effects model (GLMM) was tested to account for individual variability through random intercepts, but this resulted in a singular fit due to over-specified random effects. Therefore, a simpler GLM was used instead. The model was fitted using the glm() function in R with the binomial family. Results include coefficients (β), standard errors (SE), z-values, and p-values, with significance levels reported as “< .001”, “< .01”, or “n.s.” for non-significant effects.

Table S6.2. Detail from the model used to analyze response accuracy. The table presents the fixed effects of the model, including estimates (β), standard errors (SE), z-values, and p-values. P-values indicate significance levels, with "< .001", "< .01", "< .05", and "n.s." denoting not significant results.

| Variables | Estimate | Standard error | Z-value | P-value |
|---|---|---|---|---|
| *Intercept* | -0,264 | 0,418 | -0,632 | n.s. |
| *Block type: No-RVS* | 0,326 | 0,131 | 2,484 | < .05 |
| *Block type: Rope-RVS* | 0,197 | 0,13 | 1,516 | n.s. |
| *Block type: Snake-RVS* | 0,473 | 0,133 | 3,549 | < .001 |
| *Age* | 0,015 | 0,014 | 1,046 | n.s. |
| *Handedness: Right* | 0,018 | 0,136 | 0,135 | n.s. |
| *Sex: Male* | 0,063 | 0,094 | 0,664 | n.s |

The No-RVS condition was found to be significantly different from the Distractor condition in the glm model ($p < .05$). However, when tested with pairwise contrasts using emmeans and multiple testing correction, the difference was not significant ($p = 0.062$).
